# Platelet Factor 4 (PF4) Improves Survival in a Murine Model of Antibiotic-Susceptible and Methicillin-Resistant *Staphylococcus Aureus* Peritonitis

**DOI:** 10.1101/2023.08.25.554865

**Authors:** Nataly P. Podolnikova, Valeryi K. Lishko, Robert Roberson, Zhqian Koh, Dmitry Derkach, David Richardson, Michael Sheller, Tatiana P. Ugarova

**Author notes:** To whom correspondence should be addressed: Dr. Tatiana Ugarova, and Nataly Podolnikova.

## Abstract

The complement receptor CR3, also known as integrin Mac-1 (CD11b/CD18), is one of the major phagocytic receptors on the surface of neutrophils and macrophages. We previously demonstrated that in its protein ligands, Mac-1 binds sequences enriched in basic and hydrophobic residues and strongly disfavors negatively charged sequences. The avoidance by Mac-1 of negatively charged surfaces suggests that the bacterial wall and bacterial capsule possessing net negative electrostatic charge may repel Mac-1 and that the cationic Mac-1 ligands can overcome this evasion by acting as opsonins. Indeed, we previously showed that opsonization of Gram-negative *Escherichia coli* with several cationic peptides, including PF4 (Platelet Factor 4), strongly augmented phagocytosis by macrophages. Here, we investigated the effect of recombinant PF4 (rPF4) on phagocytosis of Gram-positive *Staphylococcus aureus* in vitro and examined its impact in a mouse model of *S. aureus* peritonitis. Characterization of the interaction of rPF4 with nonencapsulated and encapsulated *S. aureus* showed that rPF4 localizes on the bacterial surface, thus making it available for Mac-1. Furthermore, rPF4 did not have direct bactericidal and bacteriostatic activity and was not toxic to host cells. rPF4 enhanced phagocytosis of *S. aureus* bioparticles by various primary and cultured Mac-1-expressing leukocytes by several folds. It also increased phagocytosis of live nonencapsulated and encapsulated bacteria. Notably, the augmentation of phagocytosis by rPF4 did not compromise the intracellular killing of *S. aureus* by macrophages. Using a murine *S. aureus* peritonitis model, we showed that treatment of infected mice with rPF4 caused a significant increase in the clearance of antibiotic-susceptible *S. aureus* and its methicillin-resistant (MRSA) variant and markedly improved survival. These findings indicate that rPF4 binding to the bacterial surface circumvents its antiphagocytic properties, improving host defense against antibiotic-susceptible and antibiotic-resistant bacteria.

## INTRODUCTION

Platelet factor 4 (PF4) is an abundant small (7.8. kDa) cationic protein stored in mammalian platelet α-granules, accounting for ∼2% of the total granular content. PF4 is released from activated platelets at sites of vascular injury and mostly trapped inside platelet-rich blood clots where its concentration has been estimated to be 280 µM, over 100-fold greater than serum levels ^1^. The in vitro studies showed that when used in the micromolar range, PF4 can induce numerous effects on leukocytes, including migration of neutrophils and monocytes ^2^. Although PF4 has been assigned to the CXC chemokine subfamily based on its structure and chemotactic activity, this molecule does not have a typical N-terminal ELR motif required for binding to G-protein coupled chemokine receptors. The attempts to identify the receptor mediating PF4 responses were unsuccessful for several decades, and consequently, PF4’s biological function remained unclear ^1, 2^. Recently, we have identified integrin Mac-1 (CR3, CD11b/CD18) abundantly expressed on the surface of myeloid leukocytes as the primary receptor for PF4 ^3^. We showed that rPF4 supports a potent migratory response in neutrophils and macrophages, entirely dependent on Mac-1. PF4 also mediated the efficient adhesion of Mac-1-expressing cells and induced integrin clustering. We localized two segments in PF4 involved in the interaction with the α_M_I-domain, a ligand-binding domain of Mac-1, and showed that these sequences conform to the general recognition specificity of Mac-1. We previously demonstrated that in its ligands, Mac-1 binds sequences enriched in basic and hydrophobic residues and strongly disfavors negatively charged residues ^4, 5^. Like many other sequences recognized by this multiligand receptor ^5–8^, the Mac-1 binding sites in PF4, Cys^12^-Ser^26^ and Ala^57^-Ser^70^ contain several clusters of positively charged and hydrophobic residues.

Mac-1, also known as the complement receptor CR3, is one of the phagocytic receptors on the surface of neutrophils and macrophages ^9, 10^. Since the Mac-1 binding sequences in PF4 are positively charged, they can interact with the bacterial wall possessing a negative electrostatic charge. In Gram-positive bacteria such as *S. aureus*, the negative charge of the wall is due to the presence of phosphate and carboxyl groups in teichoic acids and polyanionic glycopolymers that do not have phosphate groups in their polymer backbones ^11, 12^. The positively charged PF4 sequences can also bind the bacterial capsule made of negatively charged polysaccharides ^13–16^. Indeed, the interaction of PF4 with various bacteria has been documented ^17^. Therefore, PF4 should be an ideal opsonin to bridge bacteria and Mac-1 on phagocytes. In line with this idea, we have shown that PF4 induced a potent augmentation of phagocytosis of *E. coli* by macrophages ^3^. This finding strengthened the view that PF4 is an antimicrobial protein ^18^. However, the conclusion about PF4’s antimicrobial properties has been previously made based on the report by Tang et al., who showed its direct bactericidal activity against *S. aureus* at pH 5.5 ^19^. But even so, these investigators did not observe the effect of PF4 at pH 7.2, and no bactericidal activity of PF4 against *E. coli* and *S. aureus* in a phosphate buffer system (pH 7) was found in another study ^20^, a discrepancy interpreted as a consequence of different buffer and pH conditions. The idea that the primary function of PF4 is the antimicrobial activity arising from its opsonic effect has not been explored before our studies.

To gain further insight into the role of PF4 as an enhancer of the host immune system in eradicating bacterial infections and determine whether it exhibits a broad-spectrum opsonic activity, we conducted experiments to evaluate the effect of PF4 on phagocytosis of *S. aureus,* one of the most prominent Gram-positive pathogens. Using immunofluorescence, we demonstrated that PF4 localizes on the surface of unencapsulated and encapsulated *S. aureus*, thus serving as a readily available opsonin for macrophage receptor Mac-1. PF4 was not toxic for host cells and dramatically enhanced phagocytosis of *S. aureus* by mouse neutrophils and macrophages in vitro. It also reduced bacterial burden in a mouse model of *S. aureus* peritonitis and significantly improved the survival of mice infected with antibiotic-susceptible *S. aureus* and its methicillin-resistant (MRSA) variant.

## EXPERIMENTAL PROCEDURES

### Reagents

The rat mAb M1/70 and mouse mAb 44a, which recognize the mouse and human α_M_ (CD11b) integrin subunit of integrin Mac-1, respectively, were purified from conditioned media of hybridoma cells obtained from the American Type Culture Collection (ATCC, Manassas, VA) using protein A agarose. The mouse Alexa Fluor 488-conjugated anti-Ly6G directed against neutrophil-specific lymphocyte antigen 6, locus G (catalog #127625) was from BioLegend (San Diego, Ca). The secondary antibodies, Alexa Fluor 488-conjugated goat anti-mouse IgG (H+L) (catalog **#**A-10680), Alexa Fluor 633-conjugated goat anti-rabbit IgG (catalog #A-21071), and Alexa Fluor 633-conjugated goat anti-rat IgG (catalog #A-21094) were from Thermo Fisher. The polyclonal anti-PF4 antibody was raised in rabbits using recombinant PF4 as an antigen. Calcein-AM (catalog #C3100MP), NHS-Fluorescein (catalog #46410). Heparan sulfate sodium salt (catalog #7640) and 4% Brewer thioglycolate (TG) solution were from Sigma-Aldrich (St. Louis, MO). Alexa Fluor 568-conjugated phalloidin (catalog #A12380) and the LIVE/DEAD *Bac*Light Bacterial Viability kit (catalog #13152) were from Thermo Fisher.

### Expression of recombinant PF4

To produce recombinant PF4 (rPF4), cDNA encoding human PF4 ORF was cloned into the pET-15b vector (Novagen, Madison, WI) and transformed into Origami B (DE3) competent cells (Novagen, Madison, WI). rPF4 was purified from soluble fractions of cell lysates by affinity chromatography using a 5-ml HiTrap heparin-agarose column (GE Healthcare). Endotoxin-free rPF4 was prepared as previously described ^3^. rPF4 was iodinated with ^125^I using IODO-GEN (Pierce, Rockford, IL).

### Mice

C57BL/6 and Mac-1^−/−^ (B6.129S4-Itgam^tm1Myd^/J) mice were purchased from The Jackson Laboratory (Bar Harbor, ME). All procedures were performed under the animal protocols approved by the Institutional Animal Care and Use Committee of Arizona State University. Animals were maintained under constant temperature (22 °C) and humidity on a 12-h light/dark cycle in the Animal Facility of Arizona State University. Eight- to 12-week-old male and female mice were used in all experiments with age- and sex-matched mice selected for side-by-side comparison.

### Bacterial strains and growth conditions

The antibiotic-susceptible strain of *S. aureus* subsp. *aureus* Rosenbach (catalog #25923) and methicillin-resistant *S*. *aureus* subsp. *aureus* Rosenbach strain (catalog #33591) were obtained from ATCC. Bacteria were grown overnight at 37 °C on Luria-Bertani (LB) agar plates. A single colony was transferred to LB media and incubated with shaking in screwcap 15-ml plastic tubes containing 3 ml medium for 16 h at 37 °C. The culture was diluted 100 times in fresh media and grown at 37 °C until the mid-log phase. The bacteria were harvested by centrifugation at 4000xg for 20 min, washed with PBS and adjusted to desirable CFU/ml. Colony forming units (CFU) per milliliter values were determined by colony counts of serial dilutions on agar plates incubated overnight at 37 °C. To produce encapsulated *S. aureus*, bacteria were grown under conditions that enhanced capsule production. Briefly, a single colony from an LB agar plate was grown in 10 ml LB media in a shaker incubator under aerobic conditions for 20 h at 37 °C to reach the post-exponential growth phase and then harvested and adjusted to desirable CFU/ml.

### Cells

Resident peritoneal macrophages were obtained from mice by lavage using cold PBS containing 5 mM EDTA as described ^21^. Peritoneal lavage containing neutrophils and monocytes was isolated from mice 4 h and 3 days after intraperitoneal injection of 0.5 mL of a 4% TG solution. Inflammatory macrophages from a 3-day peritoneum were isolated using the EasySep Mouse selection kit (StemCell Technologies, Vancouver, BC, Canada) with mAb F4/80 conjugated to PE according to the manufacturer’s protocol. The IC-21 murine macrophage cell line and human monocytic U937 cells were obtained from ATCC and grown in RPMI containing 10% FBS and antibiotics. The HL-60 human leukemia cells (ATCC) were cultured in IMDM (Iscove’s Modified Dulbecco’s medium, Invitrogen) supplemented with 10% FBS and antibiotics. The cells were differentiated into granulocytes by culturing in the same medium containing 1.3% DMSO for 5 days ^22, 23^. Human embryonic kidney (HEK) 293 cells stably expressing Mac-1 were previously described ^24^.

### ELISA

96-well plates (Immulon 4HBX) were coated with different concentrations of heparan sulfate in PBS for 16 h at 4 °C, washed with PBS, and different concentrations of rPF4 (0-40 μg/ml) were added to the wells for 3 h at 37 °C. After washing, rabbit polyclonal antibodies (5 μg/ml) were added for 2 h at 37 °C. The microtiter plates were incubated with goat anti-rabbit IgG conjugated to alkaline phosphatase, and the binding was detected by reaction with *p*-nitrophenyl phosphate, measuring the absorbance at 405 nm. Background binding to heparan sulfate was subtracted.

### Adhesion assay

96-well plates (Immulon 4HBX) were coated with different concentrations of heparan sulfate in PBS for 16 h at 4 °C and blocked with 1% polyvinylpyrrolidone (PVP) for 1 h at 37 °C. The cells were labeled with 7.5 μM calcein-AM for 30 min at 37 °C. The labeled cells were washed with Hanks’ balanced salt solution (HBSS) containing 0.1% BSA and resuspended in the same buffer at a 5 × 10^5^/ml concentration. Aliquots (100 μl) of labeled cells were added to each well and incubated for 30 min at 37 °C. Nonadherent cells were removed with two washes of PBS, and fluorescence was measured using a fluorescence plate reader (Perceptive Biosystems, Framingham, MA). To test the effect of PF4 on cell adhesion, different concentrations of rPF4 were added to the wells pre-coated with 10 μg/ml heparan sulfate and incubated for 3 h at 37 °C.

### Phagocytosis assays

Phagocytosis assays with adherent resident and inflammatory mouse macrophages and the mouse IC-21 macrophages were performed using *S. aureus* bioparticles conjugated to a pH-sensitive pHrodo dye (Invitrogen, catalog #A10010). Bioparticles (100 μg/ml) were incubated with different concentrations of rPF4 (10-40 µg/ml) for 1 h at 37 °C. Cells were resuspended in DMEM+10% FBS and cultured in Costar 48-well plates (2.5 x10^5^/well) for 3-5 h at 37 °C. After the medium was aspirated, adherent cells were washed and incubated with 0.5 ml of *S. aureus* bioparticle suspensions supplemented with rPF4 for 1 h at 37 °C. Cells were washed thrice with 1 ml of PBS, and photographs of five fields for each well were taken using an EVOS FL Auto (Thermo Scientific, Waltham, MA). Phagocytosed bioparticles were counted using ImageJ software.

Solution-phase phagocytosis assays with pHrodo-conjugated *S. aureus* bioparticles were performed with IC-21 cells and HL-60 cells. *S. aureus* bioparticles (10 µg) were incubated with different concentrations of rPF4 for 1 h at 37 °C. Aliquots (100 μl) of IC-21 macrophages (5×10^5^) and differentiated HL-60 cells (5×10^5^) in HBSS were incubated with rPF4-treated bioparticles for 1 h at 37 °C. Ice-cold PBS (400 µl) was added to the cell mixtures, and samples were analyzed using Attune NxT flow cytometer (ThermoFisher).

### Intracellular killing assay

IC-21 cells in DMEM (5×10^5^/0.1 ml) were placed into 96-well plates and allowed to adhere for 1 h at 37 °C in 5% CO_2_. *S. aureus* was incubated with different concentrations of rPF4 (0-50 µg/ml) for 1 hour at 37 °C, and 10 µl of the mixture containing 10^6^ CFU was added to IC-21 cells and incubated for 1 hour at 37 °C. Serial dilutions of supernatants were added to agar plates and incubated overnight at 37 °C, and phagocytosis was determined by CFU counting. Cells were washed three times with DMEM to remove nonphagocytosed bacteria, and 100 μg/ml of gentamycin in DMEM was added to the wells to kill the remaining extracellular bacteria. After incubation for 1 hour at 37 °C, cells were washed and incubated with 0.2% Triton X-100 using vigorous up and down pipetting for 3 min. Serial dilutions of lysates were added to LB agar plates and incubated overnight at 37 °C to count CFUs of intracellular bacteria. In additional experiments, after gentamycin treatment, cells were incubated in DMEM for 24 and 48 h at 37 °C in 5% CO_2_ before lysing and CFU counting. In selected experiments, adherent cells were incubated in DMEM for 3 hours after gentamicin treatment and then permeabilized with 0.2% Triton X-100 for 3 min. Cells were stained with two different nucleic dyes provided in the LIVE/DEAD *Bac*Light Bacterial Viability kit according to the manufacturer’s instructions to distinguish between live and dead bacteria. Confocal images were obtained with a Leica SP8 Confocal System (Exton, PA) using a 63x/1.4 objective.

### Determination of MIC

A single colony of *S. aureus* from an agar plate was transferred to LB media and incubated with shaking in the screwcap 15-ml plastic tubes containing 3 ml of medium for 16 h at 37 °C. The culture was diluted 20 times in fresh media and grown until the mid-log phase at 37 °C. After harvesting bacteria, 10^3^ CFU was added to the wells containing progressively decreasing concentrations of selected antibiotics or PF4. Plates were incubated for 16 h at 37 °C with shaking, and the visible growth of bacteria was evaluated after overnight incubation.

### Flow Cytometry

FACS analyses were performed to assess the expression of Mac-1 on the surface of differentiated HL-60 cells and to identify neutrophils and macrophages in the lavage isolated from the inflamed mouse peritoneum. HL60 cells (0.5×10^6^) were incubated with anti-Mac-1 mAb 44a (10 µg/ml) for 30 min at 4 °C, followed by Alexa Fluor 647-conjugated secondary antibody Cells in the peritoneal lavage were washed twice, and resuspended in HBSS. Aliquots (5×10^5^ /0.1 ml) were incubated with anti-Mac-1 mAb M1/70 (10 µg/ml), followed by Alexa Fluor 647-conjugated secondary antibody and Alexa Fluor 488-conjugated anti-Ly-6G mAb (5 μg/ml). The expression of epitopes was analyzed using Attune NxT flow cytometer (ThermoFisher).

### Immunofluorescence

*S. aureus* (1×10^8^/ml) in PBS was incubated with rPF4 for 30 min at 22 °C and washed twice in PBS. Bacteria then were incubated with rabbit polyclonal anti-PF4 antibody (1:250) for 30 min at 22 °C followed by Alexa Fluor 488-conjugated secondary antibody, washed in PBS, and resuspended in 1% paraformaldehyde. Labeled bacteria were deposited on glass coverslips using Cytospin at 2000g for 5 min, and confocal images were acquired using Leica SP8 Confocal System (Exton, PA) using a 60x/1.4 and 100x/1.4 oil objectives.

### Transmission electron microscopy

*S. aureus* grown in LB media was centrifugated at 4000g, washed twice in PBS, and resuspended in 2% glutaraldehyde. Bacteria were deposited onto carbon-coated copper grids (Electron Microscopy Sciences), stained with 2% uranyl acetate for 45 sec, and washed thrice in water. Samples were analyzed using a Talos L120C G2 microscope.

### S. aureus-induced peritonitis

8-weeks old mice were inoculated IP with 100 μl of *S. aureus* suspensions (5×10^7^ CFU) alone or *S. aureus* immediately followed by the injection of different concentrations (0-65 µg/mouse) endotoxin-free rPF4. Mice were sacrificed after 24 h, and peritoneal cavities were lavaged with 5 ml of a sterile endotoxin-free solution of PBS containing 5 mM EDTA. Aliquots of the peritoneal lavage were serially diluted and plated on LB agar plates. After 16 hours of incubation at 37 °C, colonies on the plates were counted. Data are expressed as a CFU/ml recovered from peritoneal lavage. Peritoneal inflammation was assessed by determining the total number of leukocytes in the peritoneal lavage, and differential cell counts were determined using Wright-stained cytospin preparations.

### Survival studies

Mice were inoculated I.P. with *S. aureus* (5×10^8^ CFU) without or in combination with endotoxin-free rPF4 (0.6 mg/kg). Animals were monitored for signs of mortality and morbidity every hour for the first 24 hours and then once daily for ten days. The primary endpoint used to assess the progress of infection and PF4 activity was infection-mediated death. The Kaplan-Meier survival plots were prepared to analyze the effect of PF4 on survival.

### Statistical analysis

All data are presented as the mean ± S.D. The statistical differences between the two groups were determined using a Student’s t-test. Multiple comparisons were made using ANOVA followed by Tukey’s or Dunn’s post-test using GraphPad Instat software. Differences were considered significant at *P* < 0.05.

## RESULTS

### Characterization of the PF4-S. aureus interaction

To serve as an opsonin, bacteria-bound PF4 should be available to the phagocytic receptor Mac-1 expressed on the surface of neutrophils and monocyte/macrophages. In agreement with a previous study ^17^, nonencapsulated *S. aureus*, dose-dependently bound Iodine^125^–labeled rPF4. At 10 µg/ml PF4, 0.11 fg of protein (8.5×10^6^ molecules) bound per 1 µm^2^ of bacteria. The lack of the capsule was confirmed by transmission electron microscopy (Fig. 1A). To examine further the association of PF4 with bacteria, we investigated the localization of rPF4 using confocal microscopy with polyclonal anti-PF4 antibody and secondary antibody conjugated with Alexa Fluor 488. As shown in Fig. 1B, *S. aureus* was surrounded by a layer of rPF4, indicating that PF4 accumulated on the surface. The binding of the antibody to PF4 was specific since bacteria incubated only with the secondary antibody did not show fluorescence (Fig. S1). We also examined the localization of rPF4 on the surface of *S. aureus* grown under conditions that promote capsule formation. Transmission electron microscopy confirmed the presence of a polysaccharide capsule phenotypically similar to previously reported images ^13, 25, 26^ and a thickness ranging from 115 to 610 nm in various cells (Fig. 1C). Like nonencapsulated bacteria, rPF4 formed a layer on the surface of the capsule (Fig. 1D).

**Figure 1.**
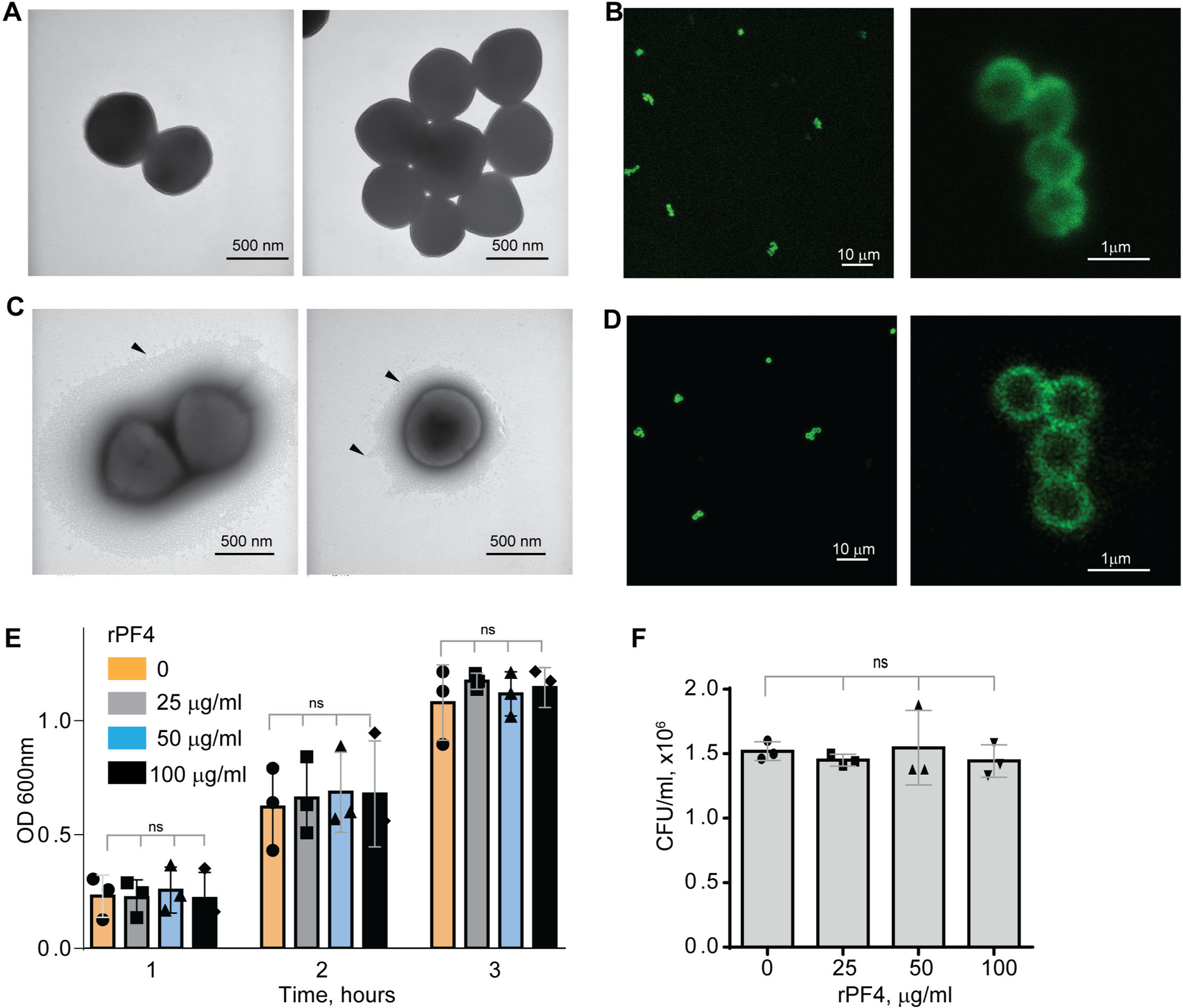
Characterization of the rPF4-*S. aureus* and rPF4-host cell interaction. Transmission electron micrographs of representative micrographs of nonencapsulated **(A)** and encapsulated **(C)** *S. aureus* stained with uranyl acetate. Arrowheads indicate the bacterial capsule in (C). The scale bar is 500 nm. **(B and D)** Confocal images showing the binding of rPF4 to *S. aureus*. After incubation with rPF4, bacteria were incubated with rabbit polyclonal anti-PF4 antibody (1:250) for 30 min at 22 °C, followed by Alexa Fluor 488-conjugated secondary antibody. The scale bars are 10 µm (*left panels*) and 1 µm (*right panels*). **(E)** Effect of rPF4 on the growth of nonencapsulated *S. aureus*. Bacteria were grown overnight in LB media and diluted in fresh LB media to OD600 of 0.03. Bacterial suspensions (0.8 ml) were incubated with different concentrations of rPF4 (25, 50, and 100 µg/ml) for 1-3 h at 37 °C, and OD 600 was measured. Data are means ± S.D. from three individual experiments; ns, no significant difference. **(F)** Bacteria grown for 3 h in the absence or presence of rPF4 were diluted 1:400,000 and cultured on LB agar plates for 24 h at 37 °C. Colonies were enumerated, and data are expressed as CFU/ml and are means ± S.D. from three individual experiments.

Since previous studies that evaluated the ability of PF4 to kill bacteria produced conflicting results ^19, 20^, we re-examined the effect of rPF4 on bacteria in the LB medium (pH 7.0) and the MES buffer (pH 5.5). *S. aureus* was incubated with different concentrations of rPF4 for 1-3 h, and bacterial growth in suspension and on LB agar plates was monitored. As shown in Fig. 1, E and F, and Fig. S2, rPF4 did not affect *S. aureus* growth at neutral and acidic pH, even at a concentration as high as 100 µg/ml. We also confirmed the lack of bactericidal activity of rPF4 compared with several antibiotics using a minimal inhibitory concentration (MIC) assay (Fig. S3). In contrast to rPF4, all tested antibiotics, except ampicillin, were active.

### PF4 is not cytotoxic for host cells

Since many antimicrobial peptides contain amphipathic α-helices that can insert into the plasma membrane, they exert not only bactericidal but also a cytotoxic effect on host cells ^27^. PF4 has the α-helix at the C-terminus, including residues 57 through 70, and a synthetic 13-residue peptide spanning the C-terminus of PF4 can kill bacteria, presumably by disrupting their membranes ^28^. Therefore, we examined the ability of rPF4 to perturb the plasma membrane by determining the leakage of calcein dye from various host cells, including leukocytes isolated from the peritoneal lavage, cultured IC-21 macrophages, and human U937 monocytic cells. rPF4 did not cause calcein leakage even at a concentration as high as 80 µg/ml (10 µM) (Fig. 2). In contrast, the cathelicidin peptide LL-37, a potent α-helical antimicrobial peptide, dose-dependently increased calcein leakage from all cells.

**Figure 2.**
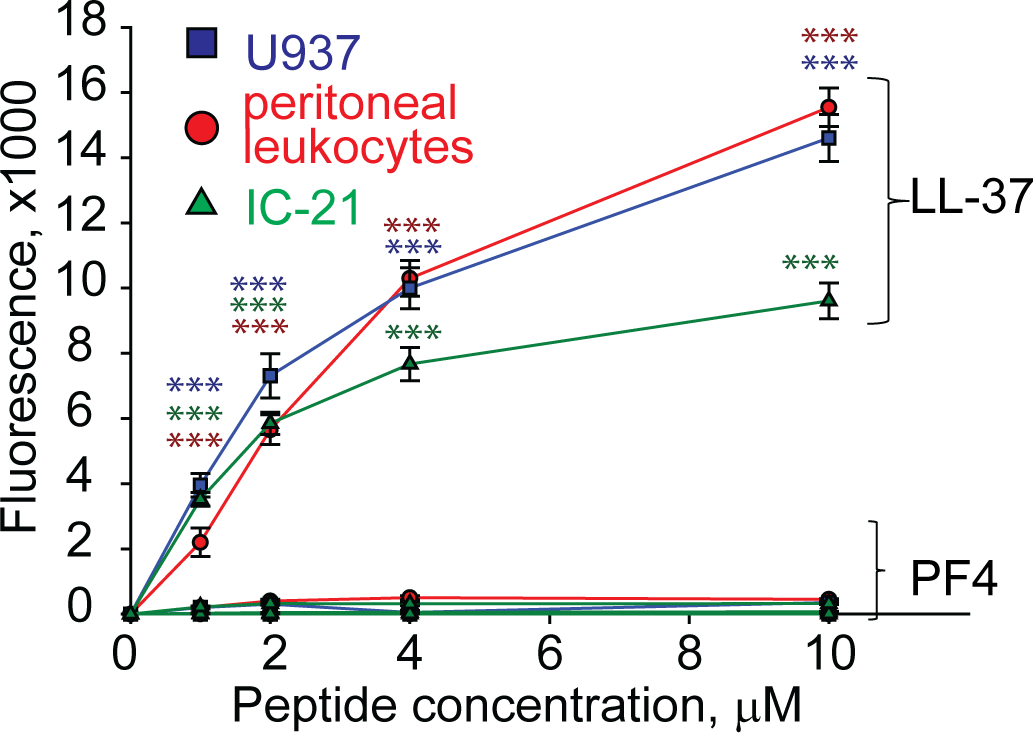
rPF4 does not have a cytotoxic effect on mammalian cells. Cultured IC-21 mouse macrophages, isolated mouse resident peritoneal leukocytes (macrophages + lymphocytes), and human U937 monocytic cells (5×10^5^/ml) were loaded with calcein for 30 min at 37 °C, washed in HBSS, and incubated with different concentrations of rPF4 and LL-37 for 30 min at 22 °C. Cells were centrifuged, and calcein leakage from cells was assessed by measuring the fluorescence of the supernatant. The effects of LL-37 and rPF4 were compared side-by-side for each cell type. The figure is representative of three experiments. The difference between LL-37 and rPF4 for each concentration is statistically significant. ***p< 0.001

### The negatively charged surfaces prevent the binding of phagocytic receptor Mac-1

The finding that Mac-1 dislikes negatively charged residues and surfaces ^4, 5^ suggests that the bacterial surface displaying negatively charged molecules may resist phagocytosis by causing the repulsion of Mac-1. To explore this possibility, in the initial experiments, we tested adhesion of Mac-1-expressing cells to surfaces coated with a negatively charged mammalian polysaccharide heparan sulfate, which is structurally similar to bacterial heparosan ^29^ and served as a model negatively-charged surface. A standard approach to test the interaction between Mac-1 and its ligands is using HEK293 cells stably transfected with Mac-1 (Mac-1-HEK293) to examine their ability to bind immobilized ligands ^4, 24^. These cells failed to attach to plastic coated with increasing concentrations (1-100 µg/ml) of heparan sulfate (Fig. 3A). Although cells attached to some extent to the uncoated plastic (∼20%), which is known to support Mac-1-mediated adhesion ^24^, only a few cells adhered to heparan sulfate. Mac-1-HEK293 cells remained round on these surfaces, a hallmark of weak adhesion (Fig. 3, D and E). Also, no adhesion of isolated peritoneal macrophages and IC-21 macrophage cell line to heparan sulfate was detected (not shown). Treatment of heparan sulfate-coated surfaces with rPF4 resulted in its binding as determined using an anti-PF4 antibody (Fig. 3B). rPF4-coated surfaces supported dose-dependent adhesion of Mac-1-HEK293 cells, IC-21, and peritoneal macrophages (Fig. 3C). Furthermore, cells could spread on rPF4-coated surfaces (Fig. 3, D and E). These data suggest that the inability of Mac-1 on macrophages to bind the negatively charged surfaces can afford bacteria protection, allowing them to evade phagocytosis, and the binding of rPF4 can remove the antiphagocytic shield.

**Figure 3.**
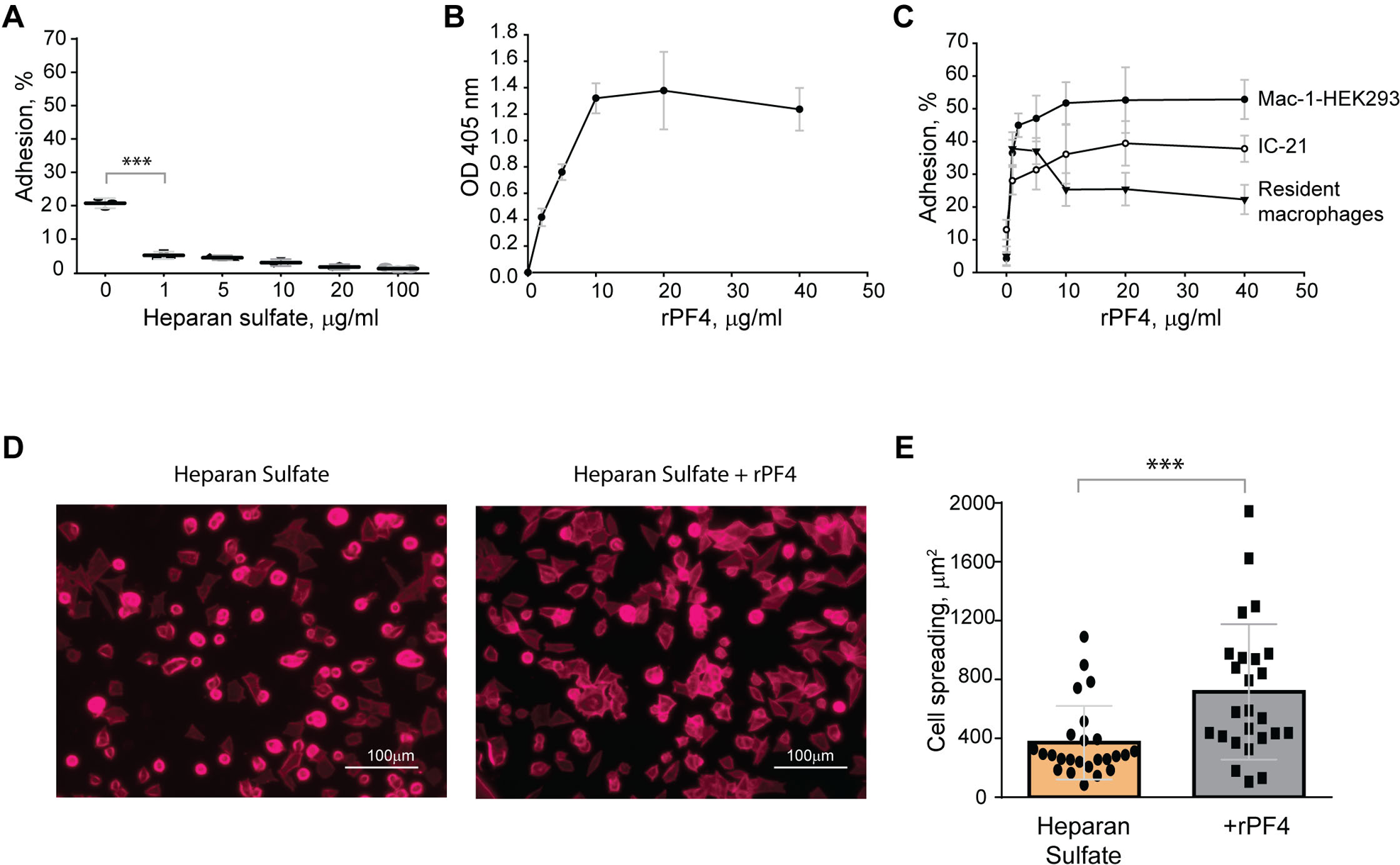
The binding of PF4 to negatively charged surfaces renders them adhesive for Mac-1-expressing macrophages. **(A)** Microtiter wells were coated with different concentrations of heparan sulfate (1-100 μg/ml) overnight at 4 °C. After washing with PBS, aliquots of calcein-labeled Mac-1-HEK293 cells (5×10^4^/0.1 ml) in HBSS were added to the wells for 30 min at 37 °C. Nonadherent cells were removed, and the fluorescence of adherent cells was measured in a fluorescence plate reader. Adhesion is expressed as the percent of fluorescence of added cells. Data are compared to cell adhesion to the uncoated plastic (0) and are means ± S.D. from three separate experiments. ***p≤ 0.001. **(B)** Microtiter wells were coated with 10 μg/ml heparan sulfate, washed with PBS, and post-coated with 1% PVP for 1 h at 22 °C. Different concentrations of rPF4 (0-40 μg/ml) were added to the wells for 3 h at 37 °C, and PF4 binding was detected using a polyclonal anti-PF4 antibody. **(C)** Microtiter wells were coated with 10 μg/ml heparan sulfate, post-coated with 1% PVP, followed by incubation with different concentrations of rPF4 (0-40 μg/ml) for 3 h at 37 °C. Aliquots (5×10^4^/0.1 ml) of calcein-labeled Mac-1-HEK293 cells, IC-21 macrophages, and resident peritoneal macrophages were added to microtiter wells. Adhesion was measured after incubation for 30 min at 37 °C. Data shown are means ± S.D. from three experiments with triplicate measurements. **(D)** Images of Mac-1-HEK293 cells spread on the surface coated with heparan sulfate (*left panel*) and heparan sulfate treated with 10 μg/ml of PF4 (*right panel*). Cells were fixed with 2% paraformaldehyde and stained with Alexa Fluor 568-conjugated phalloidin. Representative images are from three independent experiments. The scale bar is 100 μm. **(E)** Quantification of cell spreading. 25 cells from three random 20x fields were used to measure cell spreading. The cell area was determined from confocal images using ImageJ software. Data shown are means ± S.D. ***p< 0.001

### PF4 augments phagocytosis of S. aureus bioparticles by neutrophils and macrophages

To determine the effect of rPF4 on phagocytosis of *S. aureus*, we performed phagocytic assays using various populations of Mac-1-expressing leukocytes. Since the phagocytic function of neutrophils is essential for resolving microbial infections ^30^, and Mac-1 is involved in the binding of PF4 by neutrophils ^3^, we initially examined the effect of rPF4 on ingestion of *S. aureus* by HL60 cells, a neutrophil-like cell line. HL60 cells were differentiated into granulocytes (dHL-60) by culturing in DMSO, and expression of Mac-1 was verified by flow cytometry (Fig. 4A). The effect of rPF4 on phagocytosis was assessed using *S. aureus* bioparticles conjugated with a pH-sensitive pHrodo dye, which emits fluorescence only after phagocytosis in the acidic conditions of phagolysosomal compartments ^31^. Fig. 4, B and C show that rPF4 dose-dependently increased phagocytosis of *S. aureus* bioparticle by dHL-60 cells.

**Figure 4.**
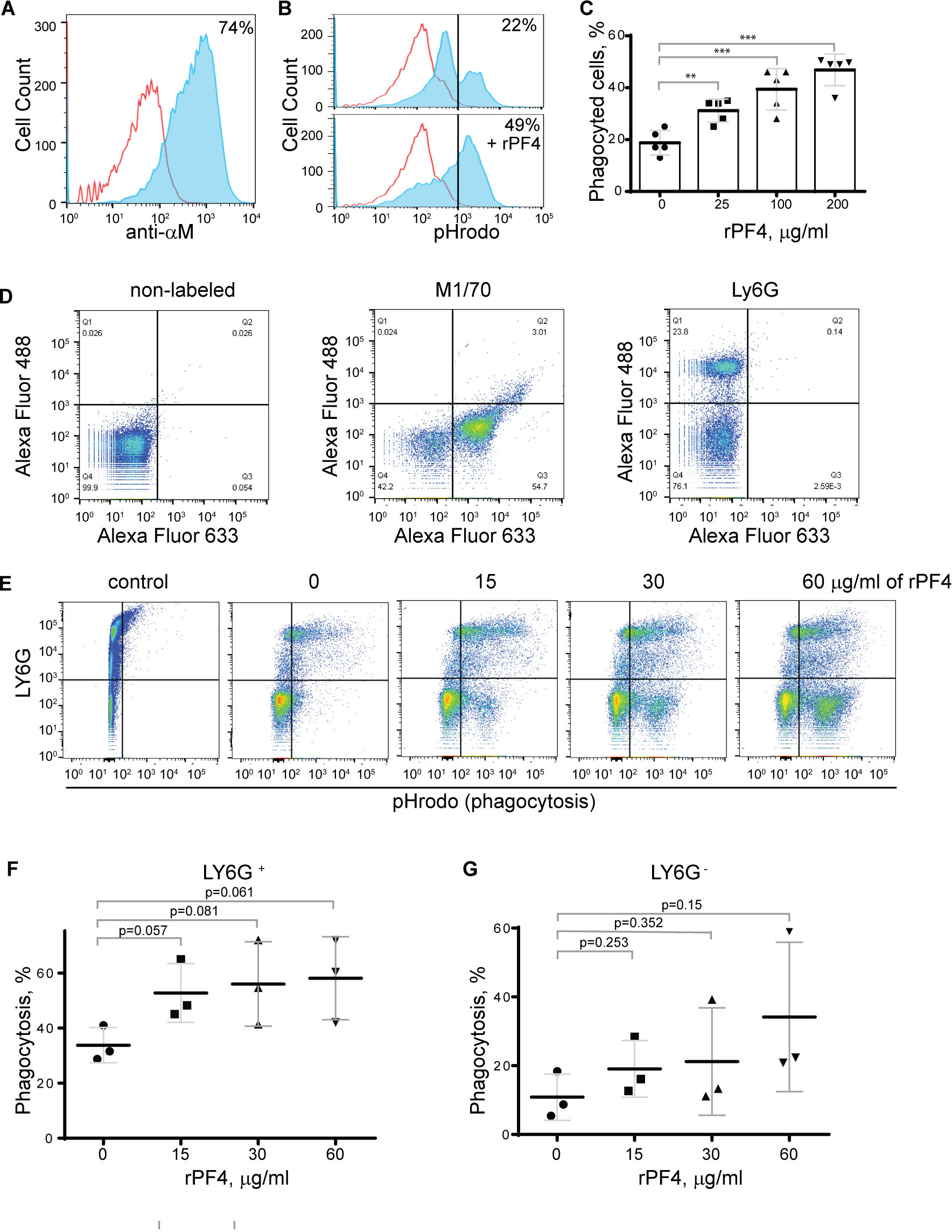
Effect of rPF4 on phagocytosis of *S. aureus* bioparticles by differentiated neutrophil-like HL-60 cells, peritoneal neutrophils, and monocyte/macrophages. **(A)** HL-60 cells were differentiated into granulocytes by incubating in the presence of DMSO, and expression of Mac-1 was determined by flow cytometry using anti-Mac-1 mAb 44a followed by Alexa Fluor 488-conjugated secondary antibody. **(B)** Flow cytometry analysis of bacteria uptake by differentiated HL-60 cells. pHrodo red-labeled *S. aureus* bioparticles (100 µg/ml) were preincubated without or with rPF4 (200 µg/ml) for 1 h and added to HL-60 cells. Phagocytosis was determined by flow cytometry after 1 h at 37 °C. Shown are representative histograms (from five individual experiments) in the absence and presence of rPF4, and numbers indicate a percentage of pHrodo-positive cells. **(C)** Effect of different concentrations of rPF4 (25-200 µg/ml) on phagocytosis of *S. aureus* bioparticles by HL-60 cells. The percentage of pHrodo-positive cells was determined by flow cytometry after 1-hour of incubation with *S. aureus* bioparticles. Values are mean ± S.D. from 5 separate experiments. **p≤ 0.01, ***p≤ 0.001. **(D)** The identification of mouse neutrophils and macrophages in peritoneum lavage 4 h after TG injection. Peritoneal cells were incubated with anti-Ly6G mAb (neutrophil marker) and anti-Mac-1 mAb M1/70 (neutrophil and macrophage marker). Shown are the total population of gated cells (*left panel*), cells expressing the M1/70 epitope (*central panel*), and cells expressing the Ly6G epitope (*right panel*). **(E)** Phagocytosis by Ly6G^+^,CD11b^+^ neutrophils and Ly6G^−^,CD11b^+^ monocyte/macrophages of pHrodo-labeled *S. aureus* bioparticles (nontreated (0) and treated with different concentrations of rPF4). Control, non-labeled cells. A representative of three separate experiments is shown. **(F, G)** Dose-dependent phagocytosis by neutrophils (Ly6G^+^) and macrophages (Ly6G^−^). Values are mean ± S.D. from three experiments; The differences in phagocytosis in the absence and presence of rPF4 are not statistically significant.

We next examined the effect of rPF4 on phagocytosis by mouse neutrophils recruited to the peritoneum by TG injection. Since both types of myeloid leukocytes, neutrophils and monocytes, migrate into the peritoneum during the first hours of sterile inflammation induced by TG injection ^32^, we used flow cytometry to distinguish the populations of cells in a 4-hour lavage. Neutrophils were identified as Ly6G^+^,CD11b^+,^ and monocyte/macrophages as Ly6G^−^,CD11b^+^ cells (Fig. 4D). The gated cell populations were then used to detect pHrodo staining in the presence of rPF4 (Fig. 4E). rPF4 increased phagocytosis of *S. aureus* bioparticles by both cell types in a dose-dependent manner (Fig. 4E). More cells in the population of Ly6G^+^,CD11b^+^ neutrophils phagocytosed *S. aureus* bioparticles in the absence of rPF4 than Ly6G^−^,CD11b^+^ monocyte/macrophages (Fig. 4F). However, although not statistically significant, Ly6G^−^,CD11b^+^ cells tended to exhibit higher phagocytic activity with increasing concentrations of rPF4 than Ly6G^+^,CD11b^+^ cells (3.1 vs. 1.7-fold) (Fig. 4G).

To further examine the effect of rPF4 on cells of the monocyte/macrophage origin, we used a mouse peritoneum-derived macrophage cell line IC-21 and peritoneal mouse resident and inflammatory macrophages. As shown in Fig. 5, A and B, rPF4 augmented phagocytosis of pHrodo-coupled *S. aureus* bioparticles by adherent IC-21 macrophages in a dose-dependent manner, and quantification indicated that at 40 µg/ml, rPF4 increased phagocytosis by ∼4-fold compared to nontreated cells. Also, at this concentration, rPF4 increased phagocytosis of bioparticles by suspended IC-21 cells by ∼5-fold (Fig. 5, C and D).

**Figure 5.**
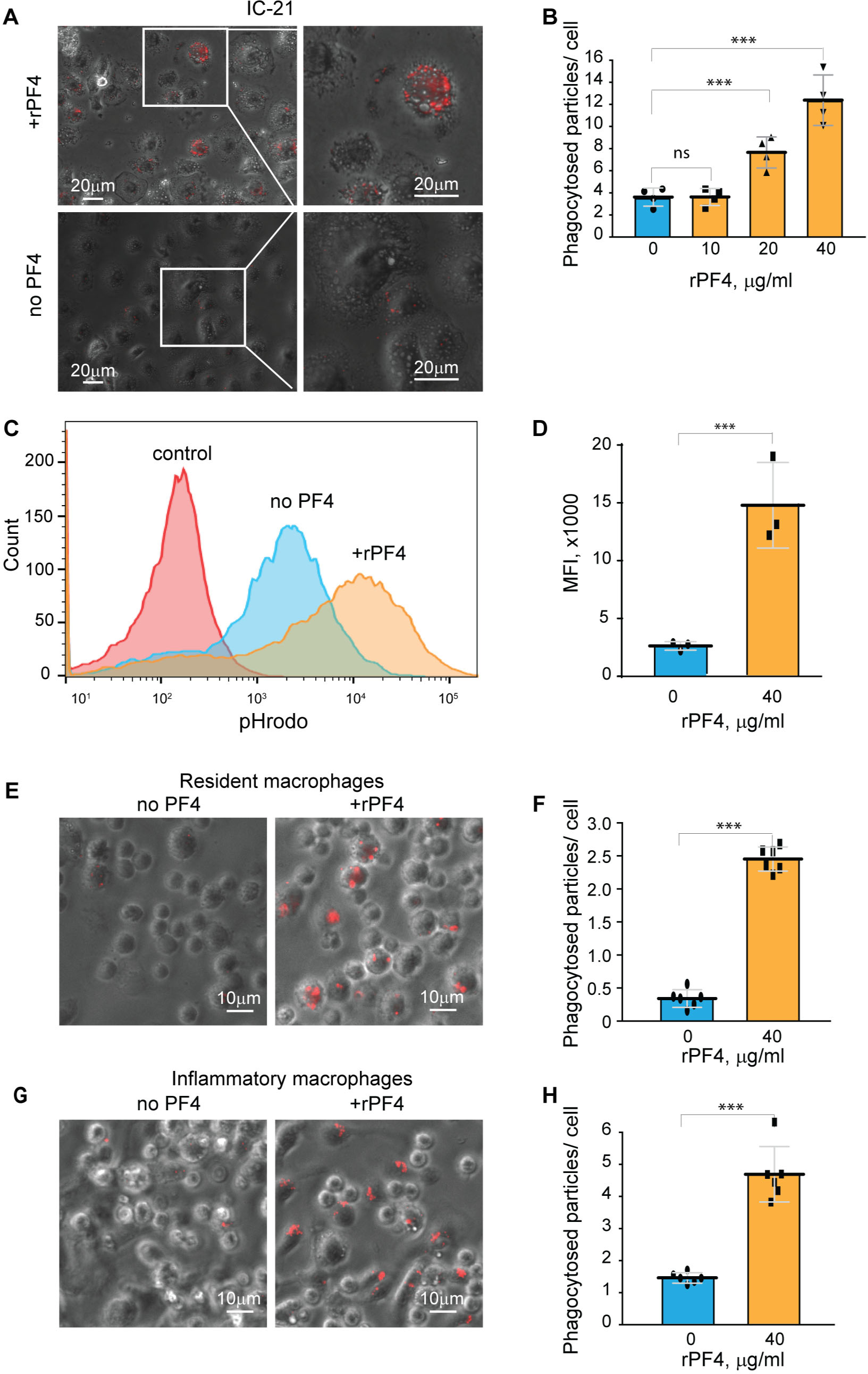
rPF4 enhances phagocytosis of *S. aureus* bioparticles by macrophages. **(A)** pHrodo-labeled *S. aureus* bioparticles (100 µg/ml) were preincubated with rPF4 (40 μg/ml) for 30 min at 37 °C, and rPF4-coated bioparticles were added to adherent IC-21 macrophages for 30 min at 37 °C. Non-phagocytosed bacteria were removed, and phagocytosis was determined from fluorescent cell images. Representative images of IC-21 macrophages exposed to rPF4-coated *S. aureus* (*upper panel*) and uncoated control bioparticles (*lower panel*) are shown. Inset boxes show enlarged regions. **(B)** Dose-dependent effect of rPF4 on phagocytosis of *S. aureus* bioparticles by IC-21 macrophages. Phagocytosis was quantified as the number of fluorescent particles per cell. Values are means ± S.D. from four random fields determined for each condition from three experiments. ***p< 0.001, ns, no significant difference. **(C)** Effect of rPF4 on phagocytosis of *S. aureus* by suspended IC-21 macrophages. Cells were incubated with pHrodo-labeled *S. aureus* bioparticles pretreated with 40 µg/ml rPF4. Phagocytosis was measured by flow cytometry. Shown are representative histograms of three experiments. **(D)** Quantification of the effect of rPF4 on phagocytosis of *S. aureus* bioparticles by suspended IC-21 macrophages from flow cytometry results. Values are means ± S.D. from three separate experiments. MFI, mean fluorescence intensity. ***p< 0.001 **(E)** pHrodo-labeled *S. aureus* bioparticles were incubated with rPF4 (40 µg/ml) and added to adherent peritoneal resident macrophages for 1 h at 37 °C. Non-phagocytosed bacteria were removed, and images of three random fields were taken. Representative images are shown. **(F)** Quantification of the effect of rPF4 on phagocytosis of *S. aureus* bioparticles by resident macrophages from results shown in (E). Phagocytosis was quantified as the number of fluorescent particles per cell. Data shown are means ± S.D. from three experiments with duplicates. ***p< 0.001 (**G, H**) Effect of rPF4 on phagocytosis of *S. aureus* by adherent inflammatory macrophages. Phagocytosis was quantified as the number of fluorescent particles per cell. Data shown are means ± S.D. from three experiments with duplicates. ***p< 0.001

Resident peritoneal leukocytes comprised of ∼60% macrophages and ∼40% B lymphocytes ^21, 33^ were obtained by lavage from the unstimulated mouse peritoneum. Recent studies demonstrated that bacterial entry into the peritoneum induces rapid macrophage adherence ^34^. Therefore, we measured phagocytosis using cells adherent to plastic after removing nonadherent B lymphocytes. At 40 µg/ml, rPF4 enhanced phagocytosis by adherent resident macrophages by ∼10-fold (Fig. 5, E and F). Since the composition and phenotype of macrophages in the peritoneal cavity change in response to infectious stimuli ^33, 35^, we also measured phagocytosis by inflammatory macrophages obtained 3 days after intraperitoneal TG injection. These cells originate from blood monocytes that infiltrate the peritoneum and gradually differentiate into the cells termed small peritoneal macrophages ^33^. Adherent inflammatory macrophages also exhibited augmented phagocytosis of *S. aureus* bioparticles in the presence of rPF4 (Fig. 5, G and H), although the effect of rPF4 was lesser than on resident macrophages. These results indicate that rPF4 significantly increases phagocytosis of *S. aureus* bioparticles by both neutrophils and macrophages.

### PF4 augments macrophage-mediated phagocytosis of live S. aureus

Since *S. aureus* bioparticles may not fully recapitulate the properties of live bacteria, we have examined the effect of rPF4 on phagocytosis of live *S. aureus*. In addition, we compared phagocytosis of nonencapsulated and encapsulated bacteria. As shown in Fig. 6A, in the absence of rPF4, resident peritoneal macrophages ingested 1.7±0.05-fold more nonencapsulated than encapsulated *S. aureus*. rPF4 enhanced phagocytosis of both strains, albeit to different extents. In particular, ∼2.8 times more encapsulated bacteria were ingested by macrophages in the presence than in the absence of rPF4 compared to ∼1.8 times for nonencapsulated bacteria.

**Figure 6.**
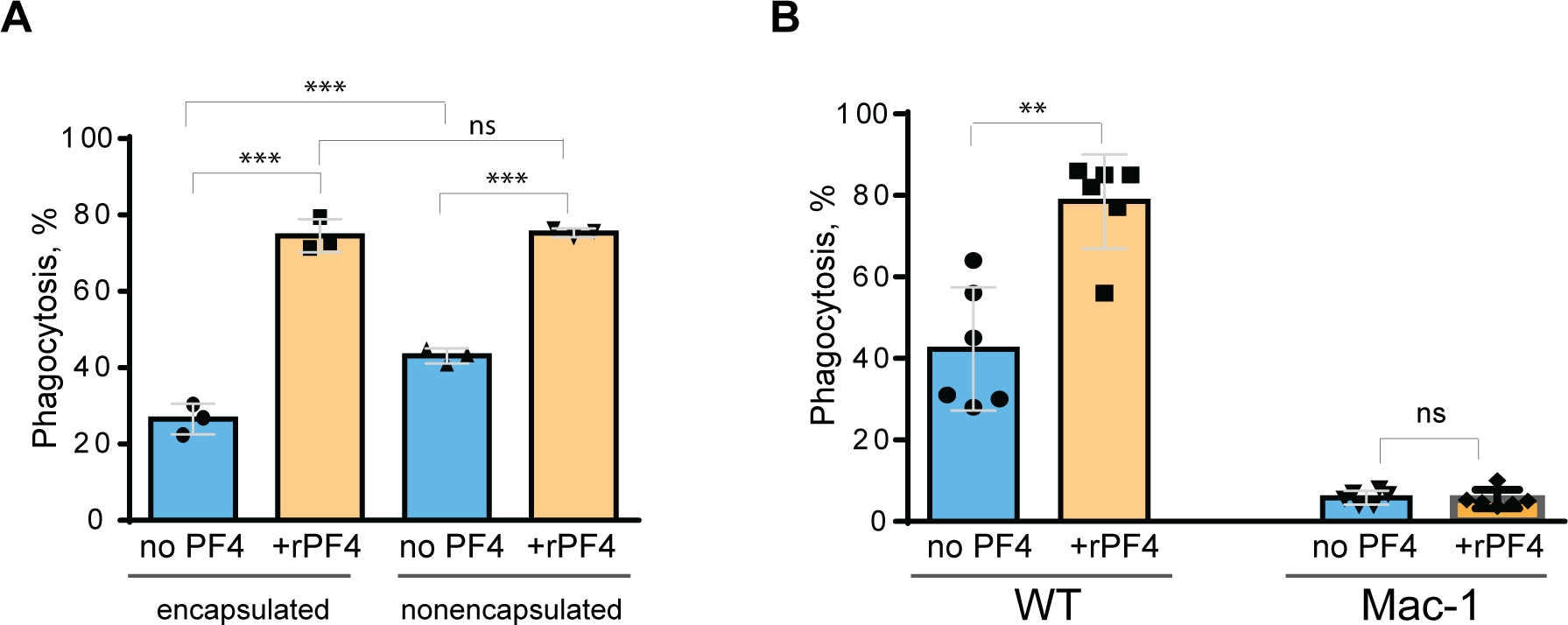
Effect of rPF4 on phagocytosis of live *S. aureus* by macrophages. Nonencapsulated and encapsulated bacteria were preincubated with 40 µg/ml rPF4 for 1 h at 37 °C. 1×10^6^ CFUs of bacteria were added to adherent inflammatory peritoneal macrophages (5×10^4^) and incubated for 1 h at 37 °C. Aliquots of media containing nonphagocytosed bacteria were plated on LB agar for 16 h at 37 °C to determine CFU/ml. Phagocytosis was assessed as a difference between total added and nonphagocytosed bacteria. Data are expressed as a percent of phagocytosed bacteria and are means ± S.D. of three experiments. ***p< 0.001, ns, no significant difference. **(B)** Adherent inflammatory macrophages isolated from WT and Mac-1-deficient mice were incubated with nonencapsulated *S. aureus* pretreated with 40 µg/ml rPF4 for 1 h at 37 °C. Data are expressed as a percent of phagocytosed bacteria and are means ± S.D. of three experiments with duplicates. **p≤ 0.01, ns, no significant difference.

We have also examined whether phagocytosis of live *S. aureus* was mediated by Mac-1. We previously demonstrated that Mac-1-deficient macrophages minimally ingested *E. coli,* and adding rPF4 did not enhance phagocytosis, indicating that the effect was specific for Mac-1 ^3^. In line with these data, only a small number of *S. aureus* was phagocytosed by Mac-1-deficient macrophages, and rPF4 did not increase their phagocytosis by these cells (Fig. 6B).

### The effect of PF4 on the intracellular killing of phagocytosed S. aureus

*Staphylococci* can survive within phagocytes ^36, 37^. To examine whether the PF4-induced augmentation of phagocytosis impacts the intracellular killing of rPF4-coated *S. aureus* in the phagolysosome, we determined the survival of internalized bacteria by quantifying CFUs of viable bacteria using the opsonophagocytic assay. Adherent IC-21 macrophages were incubated for 1 hour with nonencapsulated *S. aureus* (MOI 1:20) preincubated or not with rPF4. Counting CFUs of nonphagocytosed bacteria and subtracting this number from the initial input showed that ∼1.7- and ∼2.8-fold fewer bacteria remained in the supernatant after treatment of bacteria with 25 and 50 µg/ml rPF4, respectively, compared to control nontreated bacteria (Fig. 7A). After removing nonphagocytosed bacteria, adherent macrophages were treated with gentamicin for 1 h to remove additional extracellular bacteria, followed by cell lysis and quantitation of viable intracellular bacteria. The number of viable intracellular bacteria paralleled the augmentation of phagocytosis by rPF4 (Fig. 7B). In particular, ∼2.1 and ∼3.4-fold greater numbers were found in macrophages that ingested rPF4-treated bacteria in the presence of 25 µg/ml and 50 µg/ml, respectively, than nontreated bacteria. Also, as detected by the LIVE/DEAD BacLight assay, which allows differentiating between live and dead bacteria, an increased number of viable bacteria remained in macrophages 3 h after gentamicin treatment (Fig. 7C). However, after 48 hours, the number of colonies was drastically reduced to ≤0.1% for both treated and nontreated bacteria (Fig. 7B and Fig. S4), suggesting that the augmentation of phagocytosis by rPF4 did not compromise the ability of macrophages to carry out the effective intracellular killing. Hence, although treatment with PF4 resulted in an early increase of viable bacteria in macrophages, the numbers decreased to the same levels as for nontreated bacteria after 48 h, indicating that the intracellular killing eliminated significantly more rPF4-coated bacteria.

**Figure 7.**
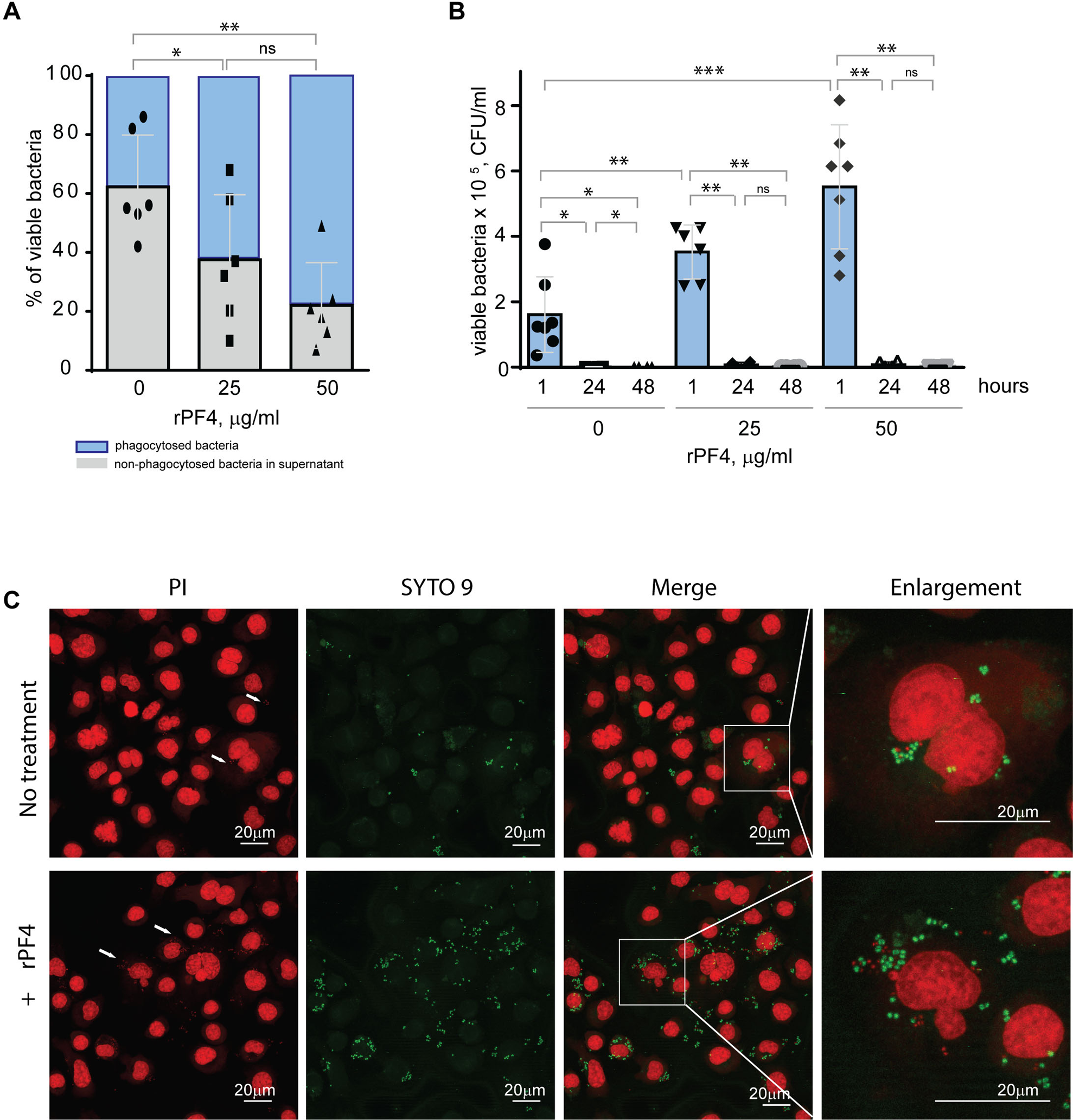
Effect of rPF4 on intracellular bacterial killing. **(A)** Adherent IC-21 macrophages (5×10^5^) were incubated with 10^6^ CFU of *S. aureus* and two concentrations of rPF4 (25 and 50 μg/ml) for 1 h at 37 °C. Aliquots of media containing nonphagocytosed bacteria were plated on LB agar for 16 h at 37 °C to count CFU/ml of viable cells. Data shown are means ± S.D. of six experiments and expressed as a percentage of nonphagocytosed (*grey*) and phagocytosed bacteria (*blue*). *p≤ 0.05, **p≤ 0.01, ns, no significant difference. **(B)** After phagocytosis, cells were washed, and extracellular bacteria potentially remaining on the cell surface were inactivated by treatment with gentamicin for 1 h, after which macrophages were either lysed with 0.2% Triton X-100 immediately or cultured in DMEM for 24 and 48 hours. Aliquots of cell lysates were plated on LB agar plates for 16 h at 37 °C, and CFUs determined. Data are means ± S.D. from six experiments. *p≤ 0.05, **p≤ 0.01, ***p< 0.001, ns, no significant difference. **(C)** Representative confocal images of macrophages stained with propidium iodide and SYTO 9 to distinguish between live and dead intracellular bacteria. Adherent IC-21 macrophages were incubated with *S. aureus* treated with 50 µg/ml rPF4 for 1 hour at 37 °C, treated with gentamicin for 1 h, and incubated for an additional 3 h in DMEM. Cells were permeabilized with 0.2% Triton X-100 and stained with a 1:1 mixture of propidium iodide (red) and SYTO 9 (green). Viable *S. aureus* bacteria are stained green, and dead bacteria are stained red. The scale bars are 20 µm.

*rPF4 enhances the clearance of S. aureus in the mouse model of bacterial peritonitis and improves survival*.

To assess the effect of rPF4 on bacterial clearance, mice were infected with 5×10^7^ CFU of *S. aureus,* followed immediately by an IP injection of rPF4. Bacterial counts in the peritoneal lavage were assessed 24 h after infection, and saline served as vehicle control. As shown in Fig. 8A, rPF4 enhanced bacteria clearance. The effect of rPF4 was dose-dependent with an IC_50_ of 5.8 ± 1.2 µg/mouse, corresponding to 0.29 ± 0.06 mg/kg. At ≥10 µg/mouse (65 µg/mouse maximal testable concentration), rPF4 reduced live bacteria in the peritoneal lavage by >90%. Analysis of cell numbers in the lavage and examination of cytospin preparations of lavage fluid from infected mice showed that whereas *S. aureus* induced a substantial influx of neutrophils and monocyte/macrophages in the peritoneum, their numbers in mice treated with 13 µg/ml rPF4 were reduced by ∼2-fold (2.3 ± 0.1 x10^6^/ml vs. 1.4 ± 0.04 x10^6^/ml for neutrophils and 2.3 ± 0.1 x10^6^/ml vs. 0.89± 0.02 x10^6^/ml for macrophages) (Fig. 8B). rPF4 injected alone slightly decreased neutrophil migration, but this difference was not statistically significant compared to the PBS control. In addition, rPF4 effectively reduced infection caused by 5×10^7^ CFU MRSA with efficacy at the same dose as for antibiotic-susceptible strain (Fig. 8C). The increased bacterial clearance in the presence of rPF4 translated into a significant difference in survival profile relative to nontreated mice. rPF4 treatment substantially improved survival from a sub-lethal dose of bacterial inoculum (5×10^8^) compared to vehicle control (Fig. 8D).

**Figure 8.**
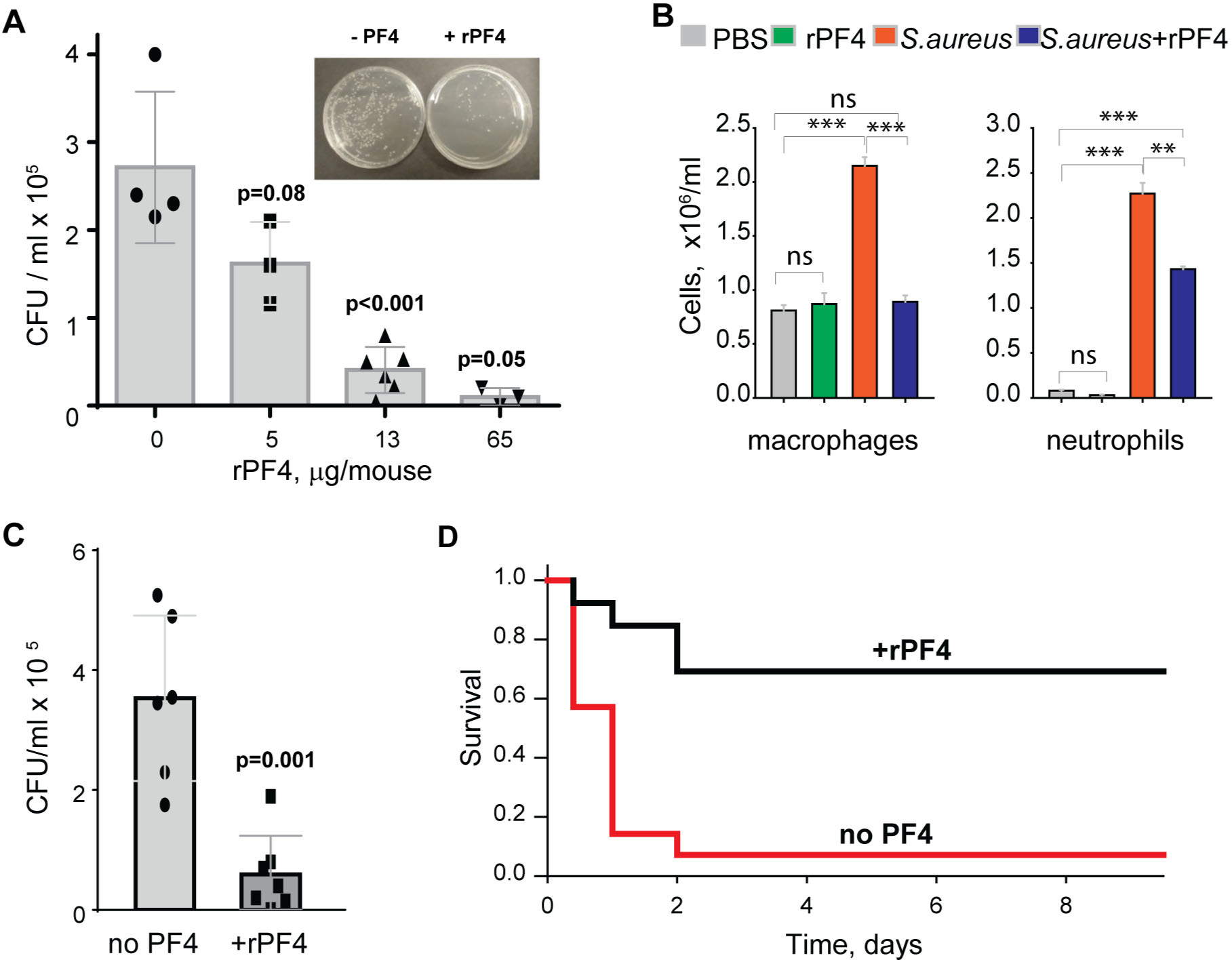
rPF4 promotes bacterial clearance and survival in a mouse peritonitis model. **(A)** The dose-dependent effect of rPF4 on bacterial clearance. C57BL/6 mice were infected IP with *S. aureus* (5×10^7^ CFU) alone or with 5, 13, or 65 μg/mouse of rPF4. After 24 hours, peritoneal lavage was collected, cells were removed by centrifugation, and the supernatant was plated on LB agar for 16 h at 37 °C. Values are CFU/ml and are means ± S.D. from 3-6 individual experiments. *Inset*, Representative images of agar plates containing bacterial colonies grown from the peritoneal lavage of control mice and mice that received 13 µg/mouse of PF4. **(B)** Differential count of cells recovered from the mouse peritoneal cavity 24 h after injection with sterile saline (PBS), rPF4 (13 μg/mouse), *S. aureus* (5×10^7^ CFU), and *S. aureus*+13 μg/mouse of rPF4. Data are means ± S.D. from four experiments. **p≤ 0.01, ***p< 0.001, ns, no significant difference. **(C)** Effect of rPF4 on clearance of MRSA. C57BL/6 mice were injected IP with MRSA (5×10^7^ CFU). The bacteria load in the peritoneal lavage was analyzed and quantitated 24 hours after infection. One group received 13 µg/mouse of rPF4 immediately after MRSA. Data are means ± S.D. derived from 7 mice in each group. **(D)** Kaplan-Meier analysis of mice (n=13 mice per group) after IP injection of *S.aureus* (5×10^8^ CFU) without or with 13 µg/mouse (0.6 mg/kg) of rPF4 reveals a significant survival benefit for PF4-injected mice. There is a statistically significant difference between survival curves, p <0.001.

## DISCUSSION

This study builds on the idea that a small cationic protein PF4 is a host defense protein performing the antimicrobial function through its opsonic activity. The opsonic effect of PF4 depends on its ability to bind to negatively charged molecules displayed on the surface of bacteria and simultaneously serve as a ligand of the phagocytic receptor CR3 (Mac-1) on neutrophils and macrophages ^3^. Our previous studies showed that, by utilizing this mechanism, rPF4 significantly enhanced phagocytosis of *E. coli* ^3^. In this study, we demonstrate that rPF4 also augments phagocytosis of *S. aureus* by myeloid leukocytes, suggesting that PF4 may possess broad-spectrum activity against Gram-negative and Gram-positive pathogens. Indeed, due to its highly cationic nature, PF4 can effectively and indiscriminately bind to the negatively charged surface of many nonencapsulated Gram-negative and Gram-positive bacteria and the bacterial capsule in encapsulated species. The characteristic feature of Mac-1 is the ability of this multiligand receptor to bind not only the complement fragment iC3b ^38, 39^ but also many cationic proteins and peptides through sequences enriched in positively charged and hydrophobic amino acid residues ^6^. We have previously identified two sequences in PF4 that serve as binding sites for Mac-1 ^3^. These sequences, ^12^CVKTTSQ<u>VRPRHI</u>TS^26^ and ^57^<u>APLYKKIIKKLL</u>ES^70^, contain cores of positively charged residues flanked by hydrophobic residues, thus conforming well to the recognition specificity of Mac-1 for cationic ligands. Therefore, the presence of two sequences that can interchangeably serve as the binding sites for bacteria and Mac-1 makes PF4 an ideal opsonin that tags bacteria, converting them into an attractive target for phagocytes.

Our in vitro studies demonstrate that rPF4 augmented ingestion of *S. aureus* bioparticles and live bacteria, nonencapsulated and encapsulated, by various Mac-1-expressing phagocytes, including cultured and primary neutrophils and macrophages. Notably, the increased uptake of PF4-treated bacteria did not compromise intracellular killing since the number of viable bacteria in macrophages after 48 h was reduced to less than 0.1%, similar to cells that uptake nontreated bacteria. In agreement with the previous report ^20^, rPF4 did not possess direct bactericidal activity at pH 7.2. Furthermore, we could not detect the antimicrobial effect of rPF4 at pH 5.5, which was reported previously ^19^. Based on these results, we suggest that the PF4-mediated protection against bacterial infection observed in our in vivo studies occurred primarily via the augmentation of phagocytosis and increased intracellular killing.

When cultivated at physiological pH, bacterial cells possess a net negative electrostatic charge ^11^. In Gram-positive bacteria, the negative charge of the wall is due to the presence of phosphate and carboxyl groups in teichoic and teichuronic acids and polyanionic glycopolymers that do not have phosphate groups in their polymer backbones ^12^. Furthermore, S-proteins, one of the most abundant bacterial surface proteins, possess a high content of acidic amino acid, giving these molecules pI values in the range from 4 to 6 ^40^. The capsule of encapsulated bacteria is also made of negatively charged polysaccharides, including polysialic acid and various GAGs ^13, 14^. We previously demonstrated that the ligand-binding α_M_I domain of Mac-1 disfavors not only the negatively charged amino acid sequences in protein ligands ^4, 6^, but, in general, negatively charged surfaces. The latter conclusion was based on our previous finding that the α_M_I domain did not bind carboxymethylated dextran covalently attached to gold coatings used in the SRP experiments ^3, 4^. In the present study, we demonstrated that Mac-1-expressing macrophages did not adhere to the surfaces coated with heparan sulfate, a model negatively charged surface, but robustly bind to PF4-treated heparan sulfate (Fig. 3). Therefore, we propose that the negatively charged bacterial wall and polysaccharide capsule protect bacteria from phagocytosis by preventing their recognition by Mac-1. Consequently, the binding of cationic PF4 to the bacterial surface results in exposure of specific Mac-1-binding sites, rendering bacteria susceptible to Mac-1-mediated engulfment and killing. The proposed mechanism of PF4 action is illustrated in Fig. 9.

**Figure 9.**
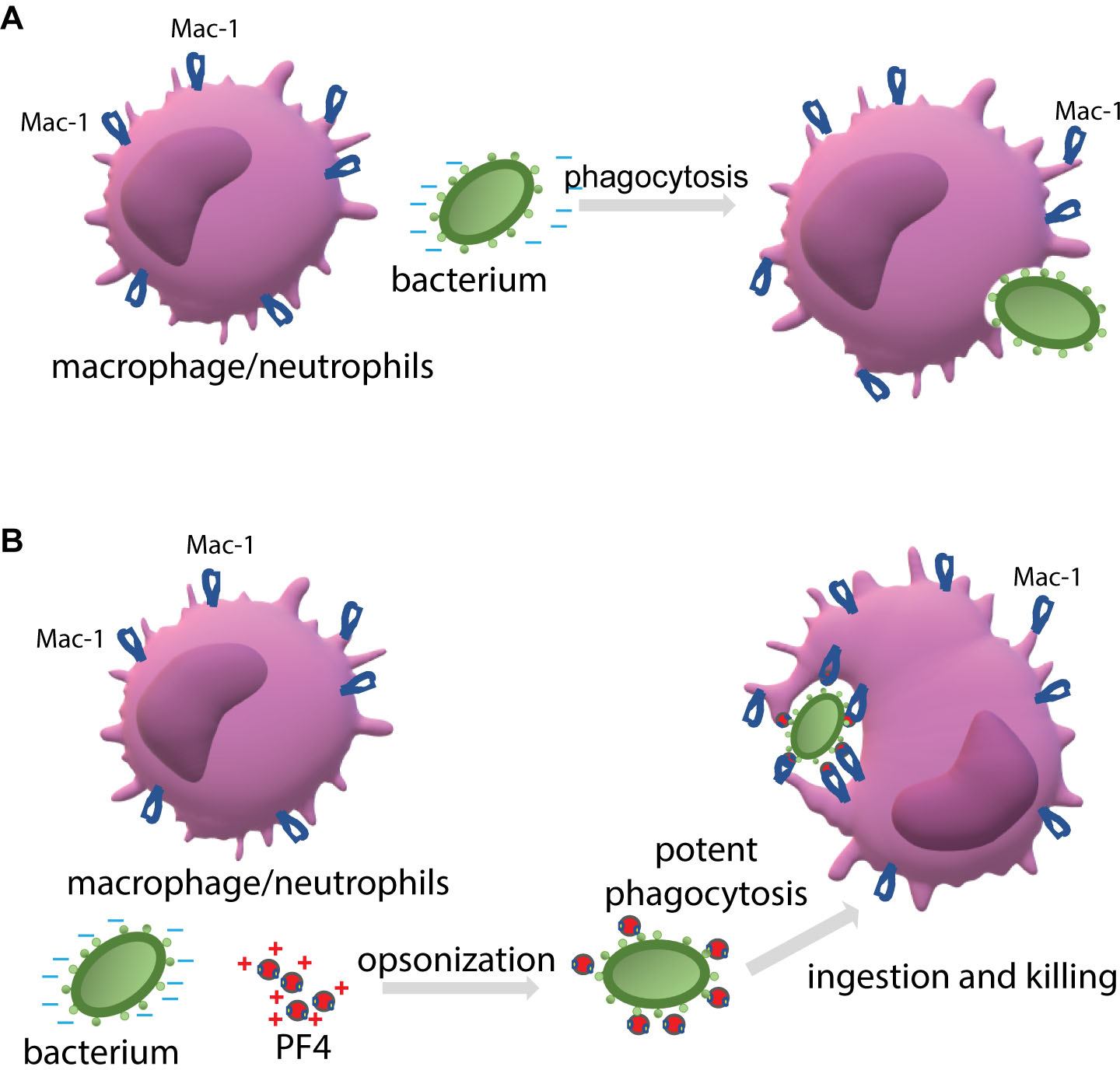
The proposed mechanism by which PF4 potentiates bacterial uptake by a phagocyte. **(A)** The phagocytic receptor Mac-1 (CR3) expressed on neutrophils and macrophages dislikes the negatively charged molecules displayed on the bacterial surface. Therefore, a bacterium evades phagocytes resulting in modest phagocytosis. **(B)** The binding of positively charged PF4, a Mac-1 ligand, to bacteria creates the binding sites for Mac-1. Thus, PF4 bridges bacteria and Mac-1, promoting the effective engulfment of bacteria by phagocytes.

It is well-established that the bacterial capsule is an important virulence factor ^13^. Previous studies correlated *S. aureus* capsule production with resistance to in vitro and in vivo phagocytic uptake and killing ^15, 41, 42, 43^. The encapsulated strains of *S. aureus* caused higher and more persistent bacteremia in mice and were more virulent in rodent models of renal infection, abscess formation, and arthritis than capsule-deficient strains ^15, 42, 44–46^. It has been proposed that the capsule resists phagocytosis due to simply repelling the negatively charged surface of phagocytes, with more highly charged capsular polysaccharides being more effective (reviewed in ^14^). Another mechanism by which bacteria avoid opsonophagocytosis is masking the components of complement and antibodies, as these opsonic proteins have been shown to deposit on the bacterial cell wall beneath the capsular layer. This steric blocking can impede their interaction with phagocytic receptors on neutrophils and macrophages, allowing the bacterium to evade phagocytic uptake ^15, 25, 47, 48^. Furthermore, the encapsulated strains bound less the complement molecules than nonencapsulated strains ^49^. The finding of PF4 as a layer surrounding the encapsulated *S. aureus* (Fig. 1) suggests that PF4 is deposited in the capsular matrix, making it available to Mac-1. Consistent with previous studies, we found that less encapsulated *S. aureus* was ingested by macrophages than nonencapsulated bacteria and treatment of encapsulated bacteria with rPF4 significantly increased the sensitivity to phagocytosis.

The property of PF4 to augment phagocytosis through its opsonic activity is not unique; other cationic antimicrobial peptides (AMPs) share this property. We and others demonstrated that the cathelicidin peptide LL-37 is a Mac-1 ligand ^7, 50^, and potent opsonin ^7^. Also, the cationic opioid peptide dynorphin A, known to exert a broad spectrum of effects on cells of the immune system, was characterized as a Mac-1 ligand and shown to enhance phagocytosis dramatically ^5^. Many other AMPs and cationic proteins from various species contain the α_M_I-domain recognition motifs, including bovine bactenecin 5, *Drosophila* drosocin, porcine cathelicidin-derived tryptophan-rich tritrpticin, synthetic peptide IDR-1 and others ^6^, and may fulfill the opsonic function. However, while the Mac-1-mediated mechanism by which LL-37, dynorphin A, and PF4 augment phagocytosis is similar, PF4 completely lacks the cytotoxic effect on host cells observed with LL-37 and dynorphin, thus distinguishing it from these molecules. Indeed, we previously showed that both LL-37 and dynorphin induced cytoplasm leakage from cultured Mac-1-expressing HEK293 cells ^5, 7^, and the present study demonstrated that LL-37 also perturbed the membrane of primary leukocytes isolated from the mouse peritoneum and cultured IC-21 macrophages. Many AMPs, including LL-37, tend to adopt an amphipathic α-helical conformation and act as membrane-active molecules that permeabilize the bacterial plasma membrane ^51^. A similar mechanism appears to be responsible for damaging host cells by LL-37 and other AMPs, causing the cytotoxic effect. Even though the PF4 segment spanning residues 57 through 70 forms the α-helix (PDB Id: 1RHP)^52^ and the synthetic peptide duplicating this sequence has antibacterial activity ^28^, we did not detect any cytotoxic effect of PF4 in concentrations as high as 80 µg/ml. The relatively high molecular weight of PF4 (∼ 8 kDa) compared to other AMPs and effective binding to the negatively charged molecules on the bacterial surface and the host cells’ glycocalyx may prevent direct contact with the plasma membrane providing a likely explanation for the lack of PF4 membrane-disruptive activities.

The key finding of the present study is that PF4 was effective in clearing bacteria in the mouse model of peritonitis and significantly improved survival. The PF4 dose required to achieve >90% bacterial clearance was ∼10 µg/mouse, i.e., ∼0.5 mg/kg. The amount of PF4 causing bacterial clearance in vivo and its concentration that enhanced phagocytosis of bacteria by macrophages in vitro (20-40 µg/ml) appear to be very similar when adjusted for the number of phagocytes (∼5×10^9^ and ∼6×10^9^ molecules per macrophage for in vivo and in vitro studies, respectively). This suggests that phagocytosis was the primary mechanism of PF4 action in the peritonitis model. Nevertheless, it remains possible that PF4 may have additional beneficial effects in vivo. In this regard, previous studies demonstrated that PF4 enhanced the generation of activated protein C (APC) ^53^, a molecule known to have anti-inflammatory effects ^54^ and prolonged survival in mice challenged with LPS dependent on the increased generation of APC ^55^. Bacteria-bound PF4 has also been shown to expose neoantigens that elicited low titers of IgG antibodies 14 days after induction of bacterial infection that potentiated phagocytosis by human neutrophils, apparently through FcγR ^17^. It is unlikely, though, that this mechanism contributed to the clearance of bacteria in our experiments because we evaluated the reduction of bacterial load after 24 h. Nevertheless, PF4 may trigger an antibody-mediated immune response against bacteria at later stages of infection, as previously suggested ^17^. PF4 has been reported to bind LPS, and this neutralizing effect may add to the action of PF4 during infections caused by Gram-negative bacteria ^56^. Finally, recent studies demonstrated that PF4, through its stabilizing effect on neutrophil extracellular traps (NETs) that are abundant in sepsis, enhanced entrapment of bacteria ^57^. Speculatively, PF4 entangled within NETs can serve as a ligand for Mac-1 on macrophages and augment bacterial phagocytosis.

Our in vitro studies demonstrate that isolated resident peritoneal and inflammatory macrophages tend to exhibit a more robust PF4-mediated phagocytic response than neutrophils. Previous studies in the peritonitis models of *S. aureus* and *E. coli* infection showed that macrophage depletion but not the elimination of neutrophils in the peritoneal cavity reduced bacterial clearance ^34, 58^, underscoring the primacy of macrophages. The underlying mechanism for the greater involvement of macrophages in PF4-mediated phagocytosis is unclear, as both macrophages and neutrophils express high levels of Mac-1. However, the higher phagocytic efficacy of macrophages compared to neutrophils in vitro suggests the mechanisms inherent to peritoneal macrophages *per se*. While the peritoneal cavity provides a good site for studying the role of PF4 in bacterial clearance because it lacks platelets, which otherwise would confound the interpretation of results due to the release of endogenous PF4, bacterial removal in this location has several specific features. In particular, clotting of peritoneal fluid in response to infection generates fibrin from fibrinogen leaking into the cavity, which can entrap bacteria aiding in the total antibacterial effect ^34, 58^. PF4 has been shown to bind fibrin ^59^, potentially creating the binding sites for bacteria that Mac-1-expressing macrophages can clear, thus assisting in phagocytosis. Further studies are required to determine whether PF4 is protective in other infection models and with other pathogens.

The global challenges drug-resistant bacteria present have stimulated the search for new direct-acting traditional antibiotics and non-traditional options ^60^. While most of these approaches have new targets or mechanisms, they still target bacteria. Only a few strategies have been proposed to promote host immune defenses. One of these host-directed strategies utilizes a small peptide SGX94 ^61^ derived from the previously characterized innate defense regulatory peptide IDR-1 ^62^, which has been shown to enhance bacterial clearance and survival in several animal infection models. The proposed mechanism of action of SGX94 is based on its ability to penetrate cells, interact directly with an intracellular protein in myeloid leukocytes and other cells, and activate signaling pathways, modulating the levels of pro- and anti-inflammatory cytokines ^61, 63^. A second strategy uses large recombinant protein plasma gelsolin (rhu-pGSN) that improves host defense by apparently stimulating macrophage NOS3 function ^64, 65^. The concentrations of these biologics shown to produce antimicrobial effects were 9.5 mg/kg for SGX9 tested in the mouse peritonitis model ^61^ and 400 mg/kg for rhu-pGSN examined in the mouse models of peritonitis induced by cecal ligation and pneumococcal pneumonia ^65, 66^. Our results show that PF4 exerted the effect at the concentration of 0.5 mg/kg (∼0.064 µmole), which, on the molar basis, is about a hundred-fold lower than SGX94 (∼8.6 µmole) or rhu-pGSN (5.1 µmole), suggesting that PF4 could be developed into an effective antimicrobial agent. Several other features make PF4 an attractive therapeutic target. One is the indiscriminate binding of PF4 to the negatively charged bacterial surfaces, which makes PF4 agnostic to specific pathogens. Notably, the finding that bacteria *per se* are entirely insensitive to PF4 and do not perceive PF4 as an enemy hints at its potential to avoid resistance ^67^.

The idea for developing PF4 into an antimicrobial agent rests not only on the elucidated mechanism of its action but also on the body of clinical observations supporting the essential role of platelets and their constituents in host defense. It is known that platelet numbers are crucial in host defense (reviewed in ^18^). For example, previous studies of experimental endocarditis demonstrated that thrombocytopenic animals had significantly higher *Streptococci* levels than their counterparts with normal platelet counts ^68^. In humans, thrombocytopenia is an independent risk factor for morbidity and mortality due to bacterial infection in elderly, oncological, and liver transplant patients and healthcare settings ^18^. In addition, inherited platelet abnormalities, including Gray platelet syndrome caused by a reduction or absence of α granules, strongly correlate with morbidity and mortality due to *S. aureus* and other infections ^69^. These findings support the idea that platelet-derived antibacterial constituents, including PF4, contribute to host defense against bacterial pathogenesis. Although PF4 is present in platelets at unusually high concentrations, thrombocytopenia, a hallmark of sepsis and other infections, may deplete the pool of PF4, resulting in the loss of its protective function. Systemic PF4 supplementation may be an efficient and controlled treatment of antibiotic-resistant bacterial infections.

## Acknowledgments

This work was supported by the NIH grant HL63199. We acknowledge the use of instruments within the Biosciences Advanced Light Microscopy Facility at Arizona State University. Confocal image data were collected using a Leica TCS SP8 LSCM (the NIH SIG award S10 OD023691). The transmission electron microscopy was performed in the Imaging Cores-Electron at the University of Arizona Research, Innovation, and Impact Core Facilities. We thank Dr. Yishin Shi, Arizona State University, for providing *S. aureus* strains and advice on growing bacteria.

## Conflict of Interest

D. Derkach and D. Richardson are employed by bioSyntagma Inc. These and all other authors declare that the research was conducted in the absence of any commercial or financial relationships that could be construed as a potential conflict of interest.

## Author contribution

Conceived and designed the analysis: **N.P**., **T.U.**

Resources: **T.U., Y.S.**

Collecting data and analysis: **N.P.**, **V.L.**, **D.D., T.U., R.R., Z.K.**

Funding acquisition: **T.U.**

Writing, Review, and Editing the paper: **T.U.**, **N.P.**, **D.D., M.S., D.R.**

## Supplemental Figures

**Figure S1. Confocal images showing specific binding of anti-PF4 antibody to rPF4-bound *S. aureus***. Nonencapsulated **(A)** and encapsulated **(B)** *S. aureus* was incubated with rPF4. After washing, bacteria were incubated with rabbit polyclonal anti-PF4 antibody (1:250) for 30 min at 22 °C, followed by Alexa Fluor 488-conjugated secondary antibody, fixed, and observed using a confocal microscope. **(C)** Control experiment showing the lack of the secondary antibody binding to the rPF4-treated nonencapsulated bacteria in the absence of the primary antibody. The scale bars are 10 μm.

**Figure S2. Effect of rPF4 on bacterial growth.** For the solution-phase microbicidal assays, *S. aureus* at 10^6^ CFU/ml were inoculated into MES buffer (pH 5.5) and incubated with different concentrations of rPF4 for 1 h at 37°C. Aliquots (100 µl) of diluted suspensions (1:500) were cultured on LB agar plates. Colonies were enumerated after incubation for 24 h at 37 °C. Data are expressed as CFU/ml and are means ± S.D. from three individual experiments. ns, no significant difference.

**Figure S3. Minimal Inhibitory concentration (MIC) of rPF4. (A)** Aliquots of *S. aureus* (10^3^ CFU) in LB were added to the wells containing serial dilutions (1:2) of rPF4 (0.4-500 µg/ml) and selected antibiotics. The visible growth of bacteria was evaluated after overnight incubation. Control, wells with no added inhibitors. **(B)** The concentrations of antibiotics or PF4 used in the MIC assay.

**Figure S4. Effect of rPF4 on intracellular bacterial killing.** Adherent IC-21 macrophages (5×10^5^) were incubated with 10^6^ CFU of *S. aureus* without or with two concentrations of rPF4 (25 and 50 μg/ml) for 1 h at 37 °C. Cells were washed, and extracellular bacteria were inactivated by treatment with gentamicin for 1 h, after which macrophages were cultured in DMEM for 24 and 48 hours and then lyzed. Aliquots of cell lysates were plated on LB agar plates for 16 h at 37 °C, and CFUs determined. The number of viable bacteria was expressed as a percent of viable bacteria present in macrophages 1 h after gentamicin treatment. Data are means ± S.D. from six experiments. ns, no significant difference.

## Notes

### Competing Interest Statement

The authors have declared no competing interest.

