## Supplemental Figures for "Platelet Factor 4 (PF4) Improves Survival in a Murine Model of Antibiotic-Susceptible and Methicillin-Resistant *Staphylococcus Aureus* Peritonitis"

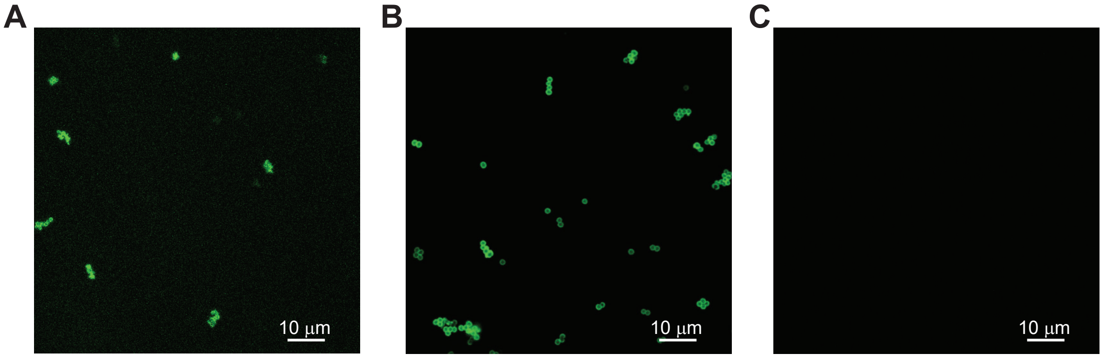


**Supplemental Figure 1. Confocal images showing specific binding of anti-PF4antibody to rPF4- bound *S. aureus*.** Nonencapsulated (A) and encapsulated (B) *S. aureus* was incubated with rPF4. After washing, bacteria were incubated with rabbit polyclonal anti-PF4 antibody (1:250) for 30 min at 22 ºC, followed by Alexa Fluor 488-conjugated secondary antibody, fixed and observed using confocal system. (C) Control experiment showing the lack of the secondary antibody binding to the rPF4-treated noncapsulated bacteria in the absence of the primary antibody. The scale bars are 10 μm.

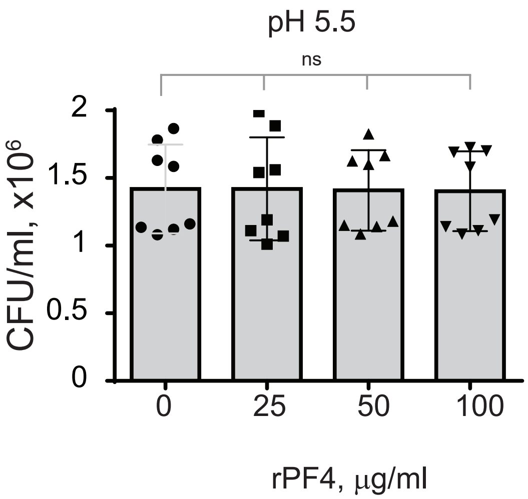


**Supplemental Figure 2. Effect of rPF4 on bacterial growth.** For the solution-phase microbicidal assays, *S. aureus* at $10^6$ CFU/ml were inoculated into MES buffer (pH 5.5) and incubated with different concentrations of rPF4 for 1 h at 37°C. Aliquots (100 μl) of diluted suspensions (1:500) were cultured on LB agar plates. Colonies were enumerated after incubation for 24 h at 37 °C. Data are expressed as CFU/ml and are means ± S.D. from three individual experiments. ns, no significant difference.

A

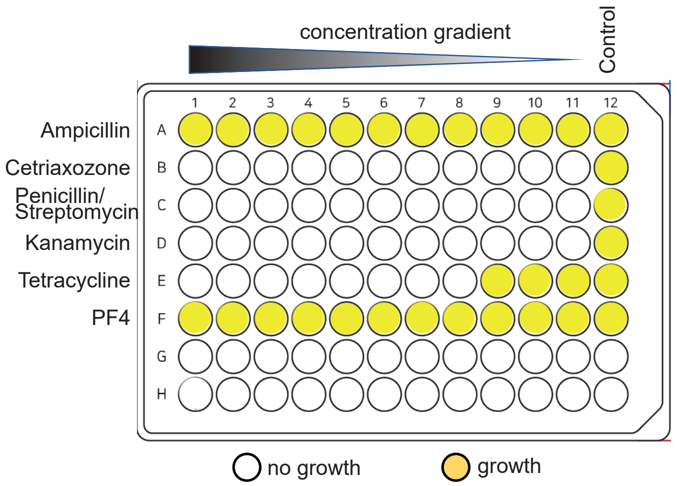


B

| Antibiotic or PF4 | Range of antibiotics, μg/ml | MIC, μg/ml |
|---|---|---|
| Ampicillin | 5-5000 | >5000 |
| Ceftriaxone | 0.02-20 | < 0.02 |
| Penicillin/Streptomycin | 2-2500 | < 2 |
| Kanamycin | 5-5000 | < 5 |
| Tetracycline | 6-6000 | 23 |
| PF4 | 0.5-500 | >500 |

**Supplemental Figure 3. Minimal Inhibitory concentration (MIC) of rPF4. (A)** Aliquots of S. aureus ($10^3$ CFU) in LB were added to the wells containing serial dilutions (1:2) of rPF4 (0.4-500 μg/ml) and selected antibiotics. The visible growth of bacteria was evaluated after overnight incubation. Control, wells with no added inhibitors. **(B)** The concentrations of antibiotics or PF4 used in the MIC assay.

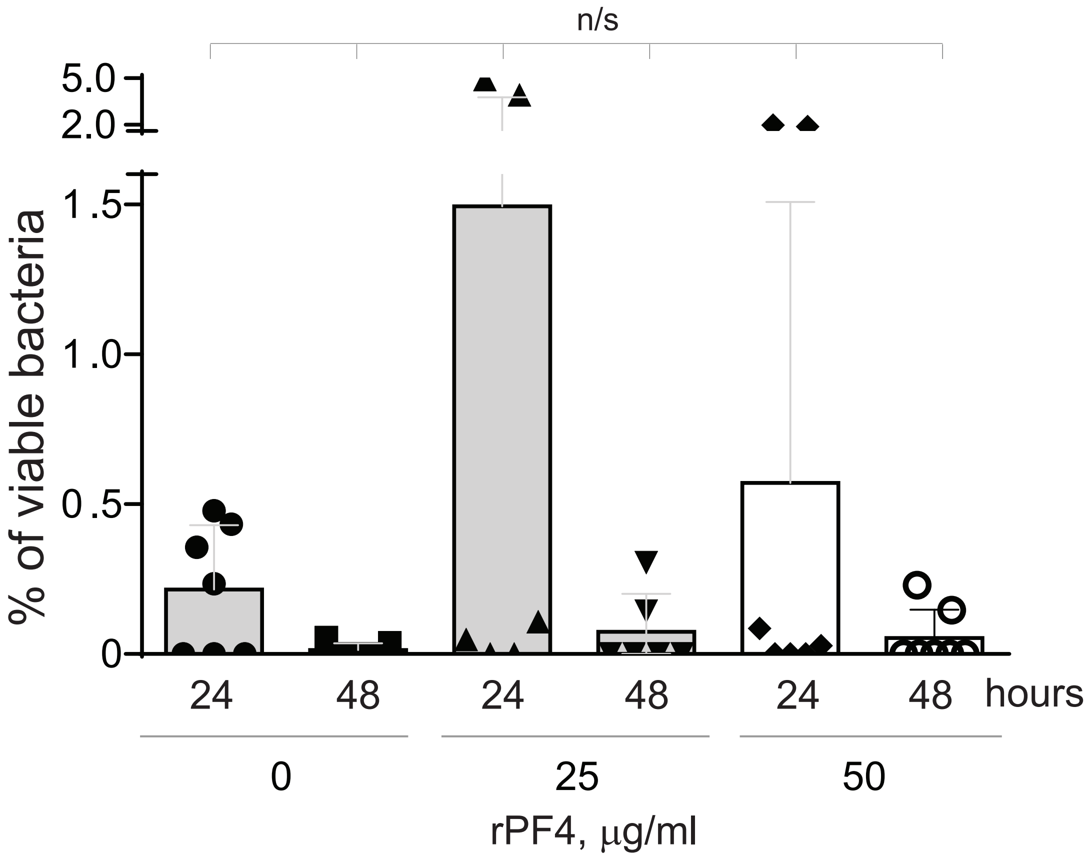


**Supplemental Figure 4. Effect of rPF4 on intracellular bacterial killing.** Adherent IC-21 macrophages (5x10$^5$) were incubated with 10$^6$ CFU of S. aureus without or with two concentrations of rPF4 (25 and 50 μg/ml) for 1 h at 37 ºC. Cells were washed, and extracellular bacteria were inactivated by treatment with gentamicin for 1 h, after which macrophages were cultured in DMEM for 24 and 48 hours and then lyzed. Aliquots of cell lysates were plated on LB agar plates for 16 h at 37 ºC, and CFUs determined. The number of viable bacteria was expressed as a percent of viable bacteria present in macrophages 1 h after gentamicin treatment. Data are means ± S.D. from six experiments.
